# Sex differences in cardio-pulmonary pathology of SARS-CoV2 infected and *Trypanosoma cruzi* co-infected mice

**DOI:** 10.1101/2021.09.18.460895

**Authors:** Dhanya Dhanyalayam, Kezia Lizardo, Neelam Oswal, Hariprasad Thangavel, Enriko Dolgov, David S. Perlin, Jyothi F. Nagajyothi

## Abstract

**Background:** Coronavirus disease-2019 (COVID-19) caused by Severe Acute Respiratory Syndrome Coronavirus 2 (SARS-CoV-2; CoV2) is a deadly contagious infectious disease. For those who survived COVID-19, post-COVID cardiac damage poses a major threat for the progression of cardiomyopathy and heart failure. Currently, the number of COVID-related cases and deaths are increasing in Latin America, where a major COVID comorbidity is Chagas’ heart disease (caused by the parasite *Trypanosoma cruzi*). Here, we investigated the effect of *T. cruzi* infection on the pathogenesis and severity of CoV2 infection and, conversely, the effect of CoV2 infection on heart pathology during coinfection.

**Methodology/findings:** We used transgenic human angiotensin-converting enzyme 2 (huACE2) mice infected with CoV2, *T. cruzi*, or coinfected with both in this study. Our study shows for the first time that white adipose tissue (WAT) serves as a reservoir for CoV2 and the persistence of CoV2 in WAT alters adipose tissue morphology and adipocyte physiology. Our data demonstrate a correlation between the loss of fat cells and the pulmonary adipogenic signaling (via adiponectin isomers) and pathology in CoV2 infection. The viral load in the lungs is inversely proportional to the viral load in WAT, which differs between male and female mice. Our findings also suggest that adiponectin-PPAR signaling may differently regulate Chagas cardiomyopathy in coinfected males and females.

**Conclusion:** We conclude that adipogenic signaling may play important roles in cardio-pulmonary pathogenesis during CoV2 infection and *T. cruzi* coinfection. The levels of adiponectin isomers differ between male and female mice during CoV2 infection and coinfection with *T. cruzi*, which may differently regulate inflammation, viral load, and pathology in the lungs of both the sexes. Our findings are in line with other clinical observations that reported that males are more susceptible to COVID-19 than females and suffer greater pulmonary damage.

## INTRODUCTION

COVID-19 illness, caused by severe acute respiratory syndrome coronavirus 2 (SARS-CoV-2; CoV2), results in debilitating disease manifestations in many infected people and increases mortality in people with comorbidities including heart diseases [1-6]. The causes of death in COVID-19 patients include cardiomyopathy, stroke, cardiac arrest, sepsis, and organ failure [7-10]. Post-COVID patients exhibit various degrees of cardiac damage, which may cause debilitating long-term effects on heart function [11-13]. Thus, the post-COVID effect may pose a major threat for the progression of cardiomyopathy and developing heart failure in patients with pre-existing heart conditions.

Although currently deaths due to COVID-19 are subsiding in many countries due to vaccination, the number of cases and deaths are still increasing in Latin America [14], where a major COVID-19 comorbidity is Chagas Disease (CD). CD is caused by the parasite *Trypanosoma cruzi*, which infects an estimated eight million people in Latin America and is also increasingly found in non-endemic countries, including 300,000 infected individuals in the United States [15]. Of these chronically infected individuals, 30% will develop chronic Chagas cardiomyopathy (CCM) and congestive heart failure, which are significant causes of morbidity and mortality [16]. Thus, vulnerable COVID-19 patients with CD are a major health burden in the Americas. In addition, the post-COVID effect on CCM in CD patients could create a health crisis in Latin America during the post-COVID era since hundreds of thousands of asymptomatic (indeterminate) CD patients likely already have or will contract COVID-19. Yet, there is virtually no clinical data or information from animal models on the interplay between CD and COVID susceptibility, severity, risk of mortality, and long-term effects on heart pathology in post-COVID CD patients.

Recent clinical meta-analysis data for COVID-19 suggest that male sex is independently associated with hospitalization, ICU admissions, need for vasopressors or endotracheal intubation and mortality [17]. Many clinical studies have also reported that male gender has been associated with a higher mortality rate due to Chagas’ heart disease [18, 19]. Male CD patients are at higher risk myocardial fibrosis and worse ventricular remodeling [19]. However, the role of sex difference in the interactions between COVID and CD is unknown.

In the present (preliminary) study, we have investigated the effect of indeterminate stage *T. cruzi* infection on the pathogenesis and severity of CoV2 infection and, conversely, the effect of CoV2 infection on heart metabolism and pathogenesis using huACE2 mice infected with CoV2, *T. cruzi*, or coinfected with both. Our results show that adipose tissue and adipogenesis play important roles in cardio-pulmonary pathogenesis during CoV2 infection and *T. cruzi* coinfection. We also demonstrate that adipogenic factors are likely responsible for (i) the observed sex-dependent susceptibility to pulmonary pathology and severity during CoV2 infection, and (ii) the pathogenesis of post-COVID cardiomyopathy or progression of post-COVID CCM in CoV2 infection or coinfection, respectively.

## MATERIALS AND METHODS

### Biosafety

All aspects of this study were approved by the Institutional Animal Care and Use and Institutional Biosafety Committee of Center for Discovery and Innovation of Hackensack University Medical Centre (IACUC 282) and adhere to the National Research Council guidelines.

### Animal model and experimental design

The transgenic mice expressing the human angiotensin-converting enzyme 2 (huACE2) (Jackson Laboratories, Bar Harbor, ME) were bred at Hackensack Meridian Health - Center for Discovery and Innovation (CDI). The Brazil strain of *T. cruzi* was maintained by passage in C3H/HeJ mice (Jackson Laboratories, Bar Harbor, ME). Both male and female mice (N=16) were intraperitoneally (i.p.) infected with 10^3^ trypomastigotes at 6 weeks of age. Mice were maintained on a 12-hour light/dark cycles and housed in groups of 3-5 per cage with unlimited access to water and chow. Once they reached indeterminate stage [20] (65 DPI; no circulating parasitemia and pro-inflammatory markers), one set of mice was coinfected intra-nasally with 1×10^4^ pfu SARS-CoV2 (NR-52281, Isolate USA-WA1/2020 COV-2 virus, NIH-BEI resources). After 10 DPI CoV2 (i.e. 75 DPI *T. cruzi* infection), we collected samples (heart, lungs, white adipose tissue (WAT) and blood; n=4/sex/subset). Age and sex matched huACE2 mice infected with SARS-CoV2 alone, as well as uninfected huACE2 mice, served as controls (Supplemental Fig. 1).

### Immunoblot analysis

Tissue lysates were prepared as previously described [20]. Each sample containing 30 μg of protein were resolved on SDS-PAGE and separately on native gel electrophoresis and the proteins were transferred to nitrocellulose membrane for immunoblot analysis. Adiponectin-specific mouse monoclonal antibody (#ab22554, Abcam), AdipoR1-specific rabbit polyclonal antibody (#ab70362, Abcam), AdipoR2-specific rabbit polyclonal antibody (#ABT12, Sigma-Aldrich), PPARα-specific rabbit polyclonal antibody (#PA1-822A, Thermo Fisher Scientific), PPARγ-specific rabbit polyclonal antibody (#2492, Cell Signaling Technology), pAMPK-specific rabbit monoclonal antibody (#2535S, Cell Signaling Technology), Cytochrome C-specific rabbit monoclonal antibody (#4280S, Cell Signaling Technology), Superoxide dismutase 1-specific mouse monoclonal antibody (#4266S, Cell Signaling Technology), Hexokinase 2-specific rabbit monoclonal antibody (#2867S, Cell Signaling Technology), β1 adrenergic receptor-specific rabbit polyclonal antibody (#12271S, Cell Signaling Technology), F4/80-specific rat monoclonal antibody (#NB 600-404, Novus Biologicals), TNFα-specific rabbit polyclonal antibody (#ab6671, Abcam), pHSL (Ser563)-specific rabbit monoclonal antibody(#4139, Cell Signaling Technology), ATGL-specific rabbit monoclonal antibody(#30A4, Cell Signaling Technology, Perilipin-specific rabbit monoclonal antibody (#D1D8, Cell Signaling Technology), IFNγ-specific rabbit monoclonal antibody (#EPR1108, Abcam), CD4-specific rabbit polyclonal antibody (#NBP1-19371, Novus biologicals), CD8-specific rabbit polyclonal antibody(#NBP2-29475, Novus biologicals), T-cadherin-specific rabbit polyclonal antibody (#ABT121, Millipore), FABP4-specific rabbit monoclonal antibody (#3544, Cell Signaling Technology), IL6-specific mouse monoclonal antibody (#66146-1-lg, Proteintech), IL10-specific rabbit polyclonal antibody (#20850-1-AP, Proteintech), BNIP3-specific rabbit monoclonal antibody (#44060, Cell Signaling Technology), Caspase 3-specific rabbit polyclonal antibody (#9662, Cell Signaling Technology) were used as primary antibodies. Horseradish peroxidase (HRP)-conjugated anti-mouse immunoglobulin (#7076, Cell Signaling Technology) or HRP-conjugated anti-rabbit immunoglobulin (#7074, Cell Signaling Technology) antibody was used to detect specific protein bands (as shown in the figure legends) using a chemiluminescence system. β-actin-specific rabbit monoclonal antibody (#4970S, Cell Signaling Technology) or Guanosine nucleotide dissociation inhibitor (GDI) (#71-0300, Invitrogen) were used as protein loading controls.

### Determination of parasite load in the tissue

After appropriate infection, organs such as heart, lungs and WAT were collected from the mice and stored at -80^0^ C. A quantitative real time polymerase chain reaction (q-RT-PCR) was used to quantify the parasite load by using PCR SYBER Green Master Mix (Roche, Applied Science, CT) containing MgCl_2_ by employing QuantStudio 3 Real-Time PCR system (Thermo Fisher). DNA isolation, preparation of standard curve and qPCR analysis was performed as previously published [21].

### Determination of SARS-CoV-2 load in the tissue

Total RNA was isolated from lungs, heart and WAT by Trizol reagent. The number of SARS-COV-2 copies were quantified using Direct One-Step RT-qPCR Mix for SARS-CoV-2 kit (Takara Bio Inc.).

### Histological analysis

Heart, lung, and WAT tissues were harvested and fixed with neutral buffered formalin overnight and embedded in paraffin wax. Hematoxylin and eosin (H&E) and Masson’s trichrome staining were performed, and the images were captured and analyzed as previously described [22]. Four to six images per section of heart or lungs were compared in each group. Histological evidence of pathology in the lungs was classified in terms of the presence of infiltrated immune cells, lipid droplets and foamy macrophages and was graded on a 6-point scale ranging from 0 to 5.

### Morphometric analysis of the heart

The hearts were harvested immediately after sacrificing the mice. The hearts were cut 5mm above the apex in cross section through the ventricles, fixed in formaldehyde, analyzed by histological staining as described earlier [23]. Briefly, the H&E sections of the hearts were used to analyze the thickness of the left ventricular wall (LVW), right ventricular wall (RVW) and the intra-septal wall [23]. The thickness of the LVW, RVW and septal wall was measured at five different locations at a magnification of 10x [23]. The average value of the 5 measurements was calculated for each mouse.

### Statistical Analysis

Data represent means ± S.E. Data were pooled, and statistical analysis was performed using a Student’s t-test (Microsoft Excel) as appropriate and significance differences were determined as p values between < 0.05 and <0.001 as appropriate.

## RESULTS

We obtained the following results using three different murine models of infections, namely, CoV2 model (infected with SARS-CoV2); *T. cruzi* model (infected with *T. cruzi*), and coinfection model (infected with *T. cruzi* followed by SARS-CoV2 at 65DPI).

### *T. cruzi* infection differently alters ACE2 levels and CoV2 load in the lungs, heart, and adipose tissue in males and females and in CoV2 infected and coinfected huACE2 mice

ACE2 is a known receptor for the cell entry of SARS-CoV2 [24, 25]. We analyzed the effect of *T. cruzi* infection on the expression levels of ACE2 in the lungs, heart, and white adipose tissue (WAT) by Western blotting (Fig. 1A). CoV2 infection significantly increased ACE2 levels in the lungs in huACE2 mice (Fig. 1A). The levels of ACE2 were significantly higher in the lungs in both male and female (5- and 3-fold, respectively) CoV2 infected mice compared to sex matched control mice (Fig. 1A). ACE2 levels significantly increased (3.5-fold) in the lungs of *T. cruzi* infected mice and were further (2-fold) increased in mice coinfected with CoV2. We observed no difference in the levels of ACE2 in the lungs between the sexes in coinfected mice. In the hearts, in uninfected female mice, the levels of ACE2 were significantly lower (2.2-fold) compared to uninfected male mice (Fig. 1A). CoV2 infection significantly increased ACE2 levels in the hearts in male (1.7-fold) and female (2.15-fold) mice, whereas *T. cruzi* infection significantly decreased ACE2 levels in the hearts in male mice (2.5-fold) (but not in female mice) compared to sex matched uninfected control mice. However, ACE2 levels in the hearts significantly increased in coinfected male and female (3.6-fold and 7.2-fold, respectively) compared to sex matched *T. cruzi* infected mice. In WAT, the levels of ACE2 were significantly higher in both male and female (2- and 1.5-fold, respectively) CoV2 infected mice compared to sex matched control mice (Fig. 1A). WAT ACE2 levels also significantly increased (1.5-fold) in *T. cruzi* infected male mice and were further (2-fold) increased in male mice coinfected with CoV2 compared to sex matched control mice. In contrast, we observed no difference in the levels of ACE2 in WAT in *T. cruzi* infected female mice and coinfected female mice compared to sex matched control mice (Fig. 1A).

**Figure 1.**
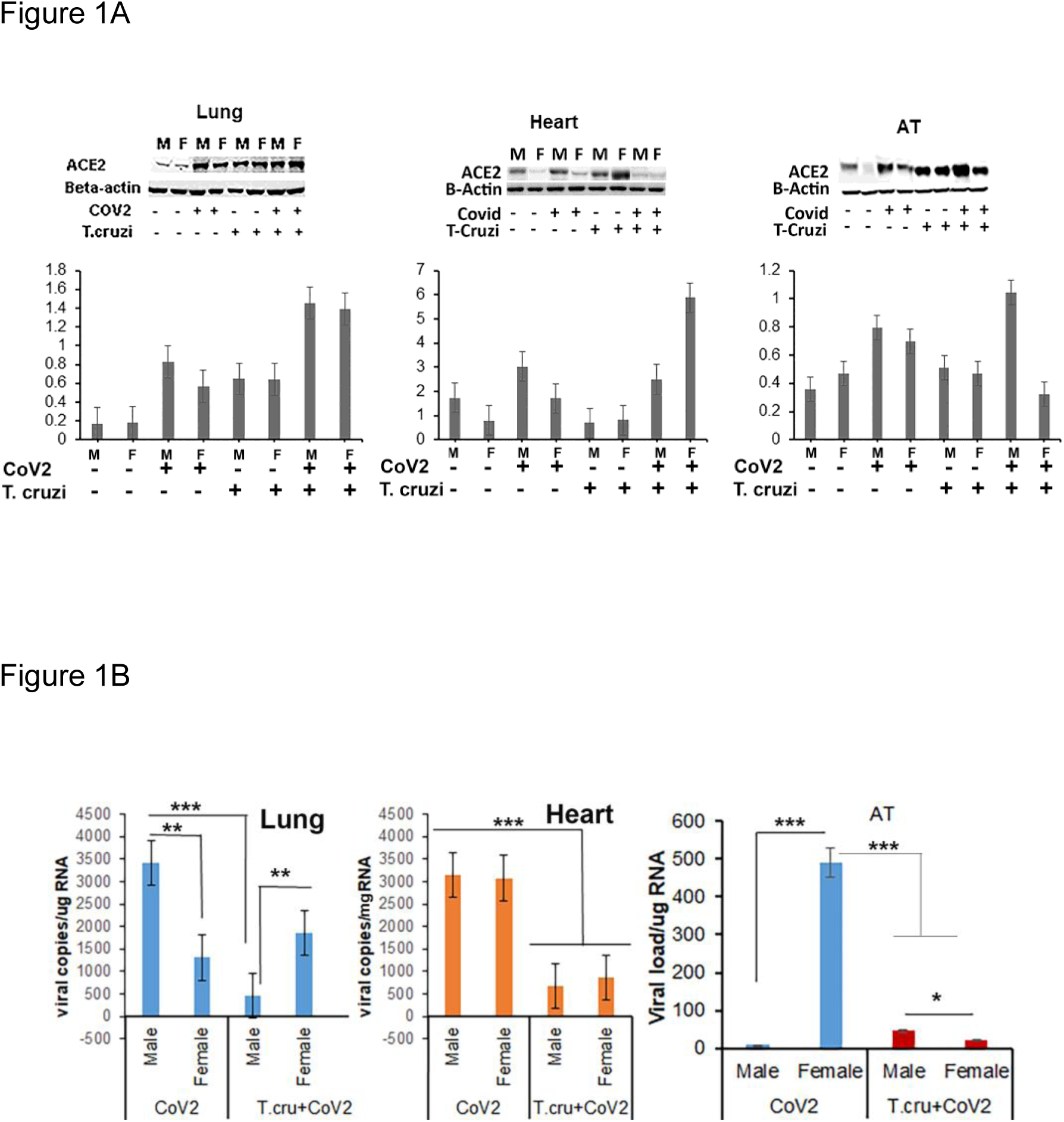
ACE2 levels and viral load in CoV2, *T. cruzi* and coinfected hACE2 mice. A) Immunoblot analysis (upper panel) of ACE2 in the lungs, heart, and adipose tissue (AT) of control and CoV2/T.cruzi and coinfected male and female mice. Bar graphs (bottom panel, x axis-arbitrary units) of the levels of ACE2 normalized to GDI. The error bars represent the standard error of the mean. The comparative-fold change in the expression levels of ACE2 are presented in Table 1A. B) Number of viral copies/ug of RNA in the lungs, and hearts, and AT quantitated by qPCR in male and female CoV2 and coinfected mice. The error bars represented the standard error of the mean (** p ≤ 0.01 and *** p ≤ 0.001). (M-male; F-female).

Lung viral loads quantitated by qPCR analysis were significantly greater in male CoV2 infected mice compared to female CoV2 infected mice (Fig. 1B), which may be due to increased ACE2 levels in male mice. Interestingly, although *T. cruzi* infection increased ACE2 levels in both male and female mice and in coinfected mice, the viral load in the lungs of female coinfected mice was significantly greater compared to male coinfected mice (Fig. 1B). However, the viral load in the lungs of coinfected male mice were significantly lower (7.5-fold, p≤0.005) compared to CoV2 infected male mice, whereas the viral load in the lungs of coinfected female mice was not significantly altered compared to CoV2 infected female mice (Fig. 1B). These data suggest that males are likely more susceptible to pulmonary CoV2 infection in general but that females may be more susceptible to pulmonary CoV2 infection in the context of CD. The-fold changes in heart ACE2 levels were significantly greater in coinfected mice compared to CoV2 alone infected mice (Fig. 1A). However, the viral load was significantly lower in the hearts of coinfected male and female mice (4.7-fold and 3.6-fold, respectively) compared to CoV2 infected male and female mice (Fig. 1B). qPCR analysis demonstrated significantly higher levels of viral load in adipose tissue in female CoV2 infected mice (64-fold, p≤0.005) compared to male CoV2 infected mice (Fig.1B). The WAT viral load in male coinfected mice was significantly higher (2-fold, p≤0.05) compared to female coinfected mice. The WAT viral load in coinfected female mice was significantly lower (23-fold, p≤0.005) compared to CoV2 infected female mice. However, the viral load in WAT in coinfected male mice was significantly higher (6-fold, p≤0.01) compared to CoV2 infected male mice. These data demonstrate that: (i) *T. cruzi* infection differently alters ACE2 levels in male and female animals; (ii) CoV2 infection differently alters ACE2 levels and viral load in male and female mice, and this difference is even greater in *T. cruzi-CoV2* coinfection; (iii) SARS-CoV2 infects and persists in adipose tissue; (iv) adipose tissue in females may act as a sink and reservoir for CoV2; and (v) an inverse relationship exists in the viral load between the lungs and adipose tissue.

### Sex difference in pulmonary pathology during CoV2 infection in *T. cruzi* infected and uninfected mice

Histological analysis of H&E and Masson’s trichrome stained lung sections of control, CoV2 infected, *T. cruzi* infected, and coinfected mice were analyzed for infiltrated immune cells, accumulated lipid droplets, fibrosis, and granulomas (Fig. 2A). Histological analysis showed significantly increased infiltrated immune cells and lipid droplets in the lungs of *T. cruzi* infected mice compared to uninfected mice (Fig. 2A and 2B). The alveolar space was more constrained and interstitial tissue thickened in male *T. cruzi* infected mice compared to female *T. cruzi* mice. CoV2 infection also significantly increased infiltration of immune cells and lipid droplets in the lungs compared to uninfected mice (sex and age matched). However, the number of granulomas and their size were greater in male CoV2 mice compared to female CoV2 mice. Interestingly, both the number and size of granulomas were greater in the lungs of female coinfected mice than in male coinfected mice. For both sexes, we observed vascular leakage (hemosiderin) and neutrophilic alveolitis in the lungs in CoV2/coinfected mice. These analyses demonstrated that: (i) The pulmonary pathology in coinfection is reduced compared to CoV2 infection alone; and (ii) although males are more susceptible to severe pulmonary CoV2 infection in general, in the context of *T. cruzi* coinfection females are more susceptible to severe pulmonary CoV2 infection compared to males.

**Figure 2.**
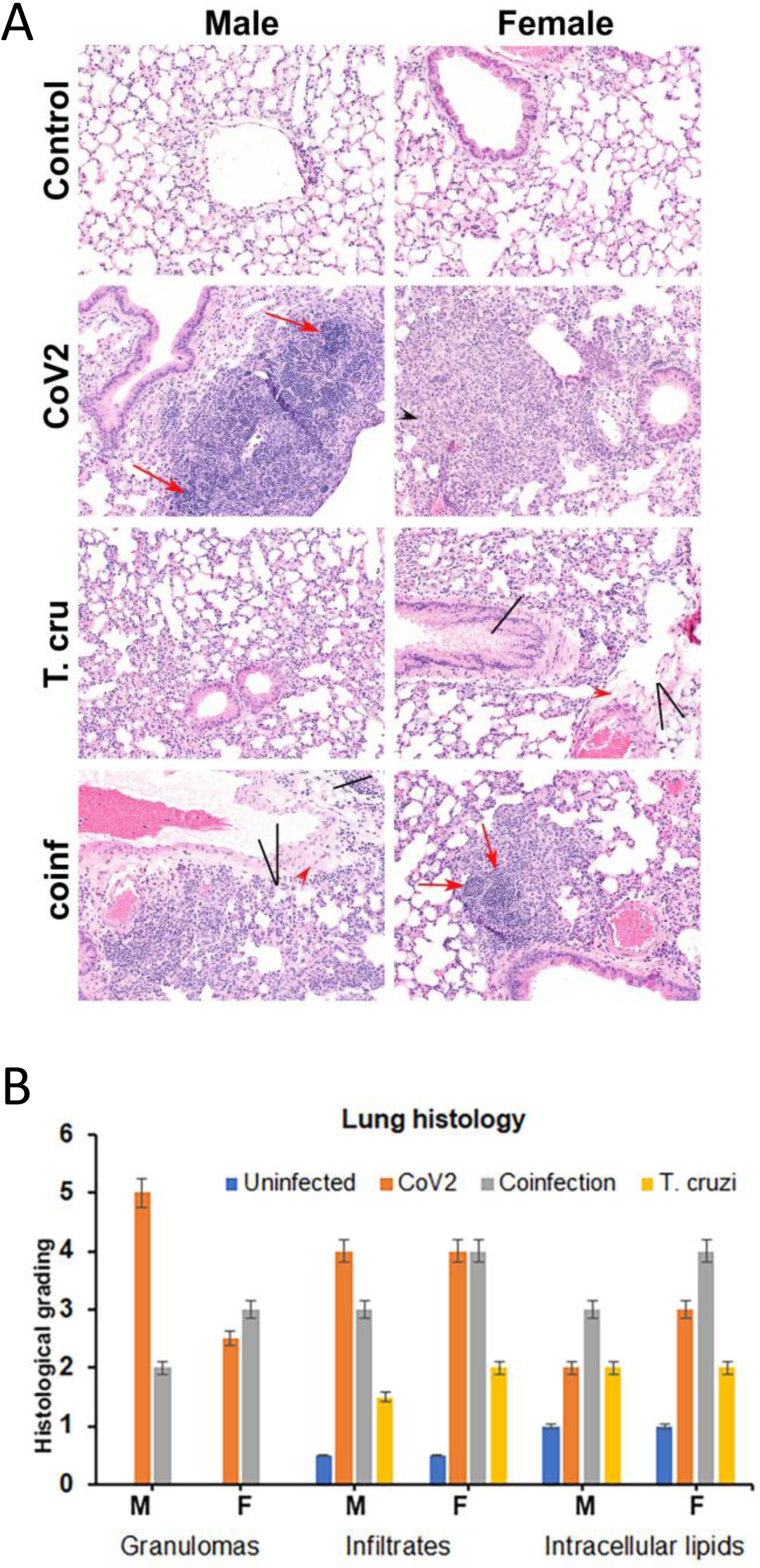
CoV2, *T. cruzi* and coinfection regulate pathology in the lungs differently in male and female hACE2 mice (n=4 mice/subset). A) Histological analysis of the lungs demonstrated increased lung pathology (infiltrated immune cells (black arrowhead) and granulomas (red arrows) and decreased alveolar space in CoV2 and coinfected mice. The presence of lipid droplets [black line] and fibrosis are shown in the images (x20 magnification). B) Histological grading of the lung’s pathology was carried out according to experimental groups and classified in terms of degree of infiltration of immune cells, granulomas and accumulation of lipid droplets. Each class was graded on a six-point scale ranging from 0 to 5+ as discussed in Method section, and presented as a bar graph.

### CoV2 infection alters adipogenic signaling in the lungs of uninfected and *T. cruzi* infected mice

Because we observed significantly increased lipid droplets in the lungs in CoV2 and *T. cruzi* infected mice compared to uninfected mice, we examined and quantified the levels of adipogenic markers such as adiponectin (ApN) and its receptors in the lungs by Western blotting. We measured the levels of lung high-molecular weight ApN (L-HMW ApN), a.k.a. its anti-inflammatory/anti-fibrotic/metabolically active form [26, 27] by native gel, and lung gAd (L-gAd), a.k.a. its pro-inflammatory form [28], by SDS-Page Western blotting (Fig. 3A). The-fold changes in the levels of HMW and gAd in the lungs in CoV2 and *T. cruzi* infected mice compared to their respective controls are shown in Table 1A. The levels of L-HMW ApN and gAd significantly increased (1.5- and 14-fold, respectively) in CoV2 infected female mice compared to uninfected female mice. However, L-HMW ApN was reduced (1.2-fold) in CoV2 infected male mice compared to uninfected male mice. *T. cruzi* infection significantly increased L-HMW ApN in both males and females (3- and 2-fold, respectively) and gAd (3.8-fold) only in female mice compared to sex matched uninfected mice. During coinfection, the levels of L-HMW ApN significantly reduced (3.6- and 2.6-fold, respectively) and gAd significantly increased (6.0- and 2.5-fold, respectively) in both male and female coinfected mice compared to sex matched *T. cruzi* infected mice. Our results suggest that CoV2 infection increases gAd levels in the lungs in *T. cruzi* infected mice.

**Figure 3.**
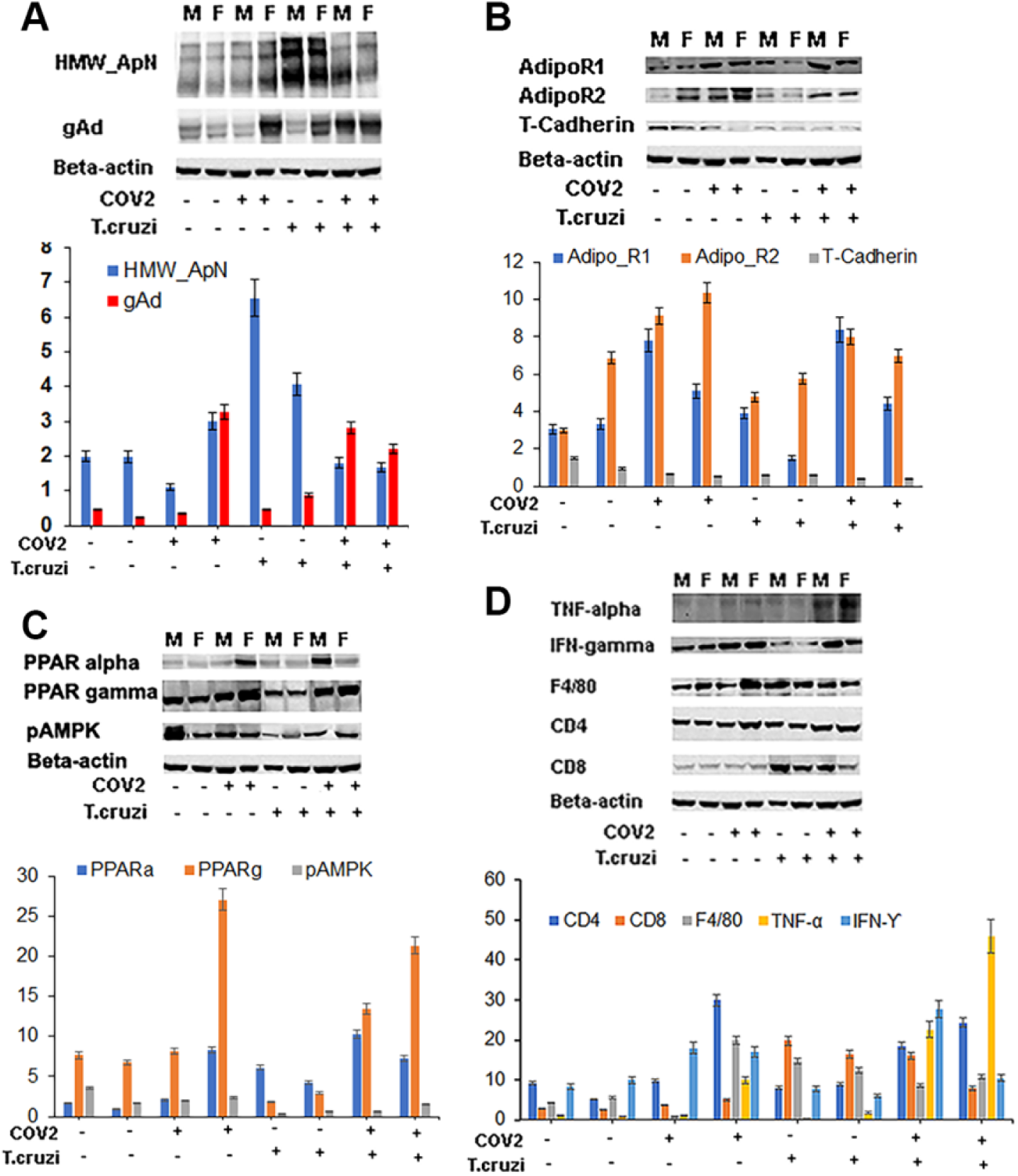
Immunoblot analysis (upper panel) of markers of :(A) adiponectin (HMW ApN and gAd); (B) ApN receptors (Adipo R1, R2 and T-cadherin; (C) lipid metabolism (PPARα and PPARγ) and energy sensor (pAMPK) and (D) Immune cells (CD4, CD8 and F4/80) and inflammatory cytokines (TNF, IFNg); in the lungs of control and infected (CoV2, *T. cruzi* and coinfected) male (M) and female (F) mice. Bar graphs (lower panels) of the levels of each protein marker normalized to either GDI or beta-actin for A, B, C, and D, respectively (x axis-arbitrary units). The error bars represent the standard error of the mean. The comparative-fold change in the expression levels of the above protein markers is presented in Table 1A.

**Table 1A.**
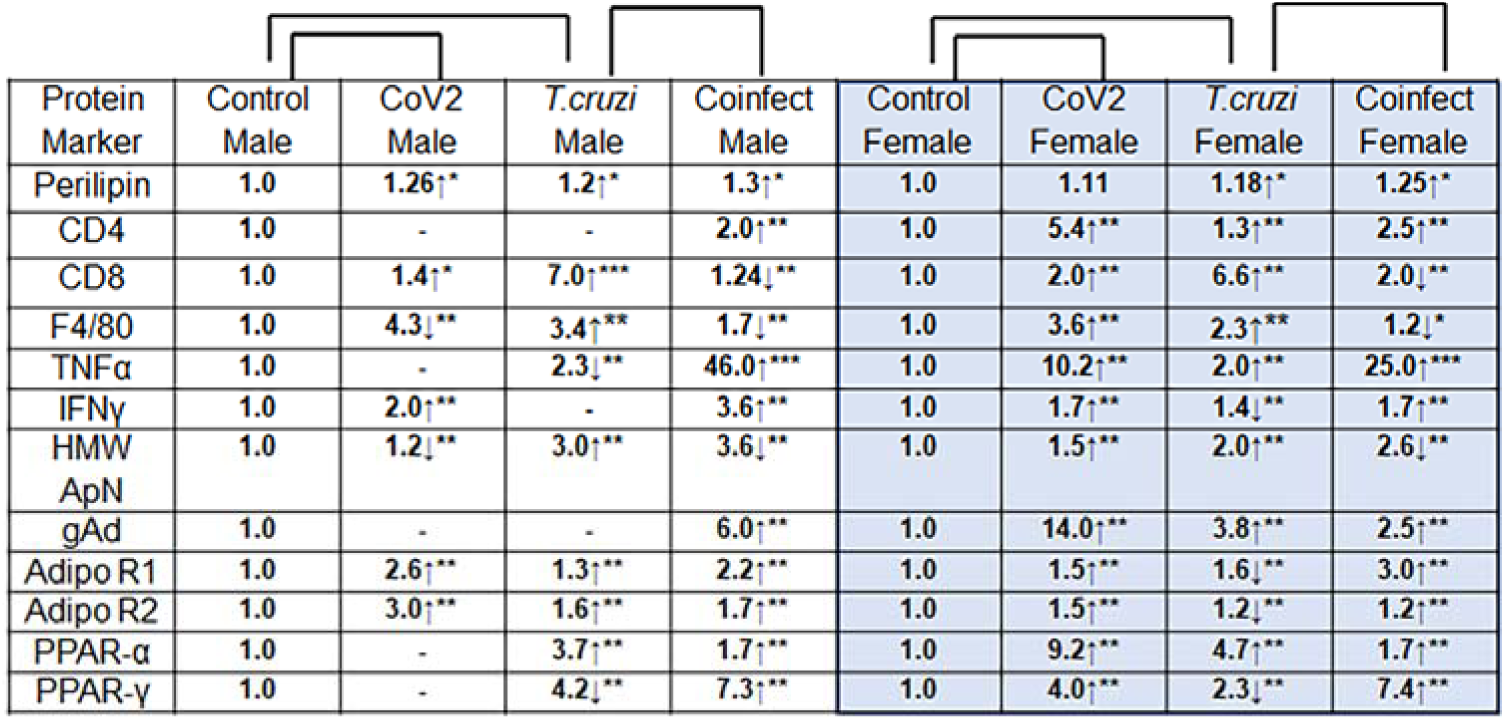
The-fold change of the protein markers (adipogenic, immune and metabolic signaling) levels compared to their sex matched control mice are presented in Table 1A analyzed in the lungs. The-fold changes were analyzed by comparing the protein’s normalized level (GDI or β-actin) in infected groups (CoV2/*T. cruzi*) to that in uninfected (control) mice, for males and females separately. For the coinfected mice, since the baseline is *T. cruzi* infection, the-fold change was calculated for coinfected mice relative to *T. cruzi* infected mice (for males and females separately). The increase and decrease in the comparative-fold change are presented by an upward or downward arrow, respectively (* p ≤ 0.05, ** p ≤ 0.01 and *** p ≤ 0.001 represents the significance). N=4/group.

The regulatory actions of ApN are mainly mediated by its receptors Adiponectin-R1 and -R2 (Adipo R1 and R2) and T-cadherin. CoV2 infection significantly increased the levels of R1 and R2 in the lungs in males (2.6- and 3.0-fold, respectively) and females (1.5- and 1.5-fold, respectively) compared to uninfected sex matched mice (Fig. 3B, Table 1A). *T. cruzi* infection significantly increased the levels of R1 and R2 in the lungs of males (1.3- and 1.6-fold, respectively) but significantly decreased both in the lungs of females (1.6- and 1.2-fold, respectively) compared to uninfected sex matched mice. Although the levels of R1 and R2 differed between male and female *T. cruzi* infected mice, R1 and R2 significantly increased in both male and female coinfected mice (Fig. 3B), suggesting that CoV2 infection increases the levels of R1 and R2 in the lungs.

ApN signaling and adipogenesis is regulated via peroxisome proliferator-activated receptors (PPARs). We analyzed the levels of PPARγ and PPARα by immunoblotting analysis (Fig. 3C). CoV2 infection significantly increased PPARγ and PPARα (4.0- and 9.2-fold, respectively) in the lungs in female mice but did not change their levels in male mice compared to sex matched uninfected mice. *T. cruzi* infection significantly decreased PPARγ but significantly increased PPARα in both males (4.2- and 3.7-fold, respectively) and females (2.3- and 4.7-fold respectively) compared to sex matched uninfected mice. However, coinfection significantly increased PPARγ and PPARα in the lungs in both males (7.3- and 1.7-fold, respectively) and females (7.4- and 1.7-fold, respectively) compared to sex matched *T. cruzi* infected mice. These data suggest that CoV2 infection induces adipogenic signaling in the lungs of female mice and male and female *T. cruzi* infected (coinfected) mice via increased PPARγ signaling.

### Immune signaling in the lungs differs between CoV2 infected and coinfected mice

Immunoblot analysis of lung lysates demonstrated significant differences in the levels of lung immune cell (CD4, CD8, and macrophage marker F4/80) and proinflammatory markers (TNFα and IFNγ) between the sexes and infections (Fig. 3D). The changes in the normalized protein levels of CD4, CD8 and F4/80 (normalized to GDI levels) are presented as a bar graph (Fig. 3D), and the relative fold change and significance are shown in Table 1A. These data showed that CoV2 infection increased CD4, CD8 and F4/80 in the lungs of female mice but increased only CD8 in male mice. *T. cruzi* infection increased CD8 and F4/80 levels in the lungs of both male and female mice and increased CD4 only in female mice. The coinfection significantly increased only CD4 levels in both male and female coinfected mice, whereas the levels of CD8 and F4/80 significantly decreased in both male and female coinfected mice compared to sex matched *T. cruzi* infected mice (although the levels of CD8 and F4/80 were higher compared to mice infected with only CoV2) (Fig. 3D).

The levels of proinflammatory TNFα and IFNγ significantly increased in the lungs in female CoV2 infected mice, with only IFNγ increasing in male CoV2 infected mice compared to sex matched uninfected mice (Table 1A). *T. cruzi* infection increased TNFα only in female mice. However, both TNFα and IFNγ significantly increased in male (46-fold and 3.6-fold; p≤ 0.005 and ≤0.01, respectively) and female (25-fold and 1.7-fold; p≤ 0.005 and ≤0.01, respectively) coinfected mice compared to sex matched *T. cruzi* infected mice (Table 1A). Our data suggest that proinflammatory TNFα is significantly higher in the lungs of female CoV2 mice and male coinfected mice compared to their sex matched infected/coinfected groups (Table 1A).

### CoV2 infection and Coinfection differently alter adipogenic signaling in WAT in male and female mice

We analyzed the levels of HMW (multimer) and LMW (trimer) adiponectin in WAT by native PAGE followed by Western blotting [29]. The levels of adiponectin (multimers and trimers) significantly increased in female CoV2 infected mice (4-fold and 24-fold, respectively) compared to sex matched control mice (Fig. 4A). The levels of multimeric adiponectin significantly decreased (2-fold) but trimers significantly increased (1.7-fold) in male CoV2 infected mice compared to male control mice. In addition, the adiponectin levels in female CoV2 mice were significantly higher compared to male CoV2 infected mice (Table 1B). On the contrary, *T. cruzi* infection significantly increased the levels of both multimer and trimer adiponectin in male mice (2-fold and 14-fold, respectively) and only trimers in female mice (2-fold) compared to sex matched control mice (Fig. 4A). Interestingly, CoV2 infection in *T. cruzi* infected mice further increased the levels of both multimer and trimer adiponectin in male mice (1.2- and 3.6-fold, respectively) and significantly reduced in female mice (2-fold and 4.7-fold) compared to sex matched *T. cruzi* infected mice, which is contrary to the changes in adiponectin levels in males and females in only CoV2 infected mice compared to control mice.

**Figure 4.**
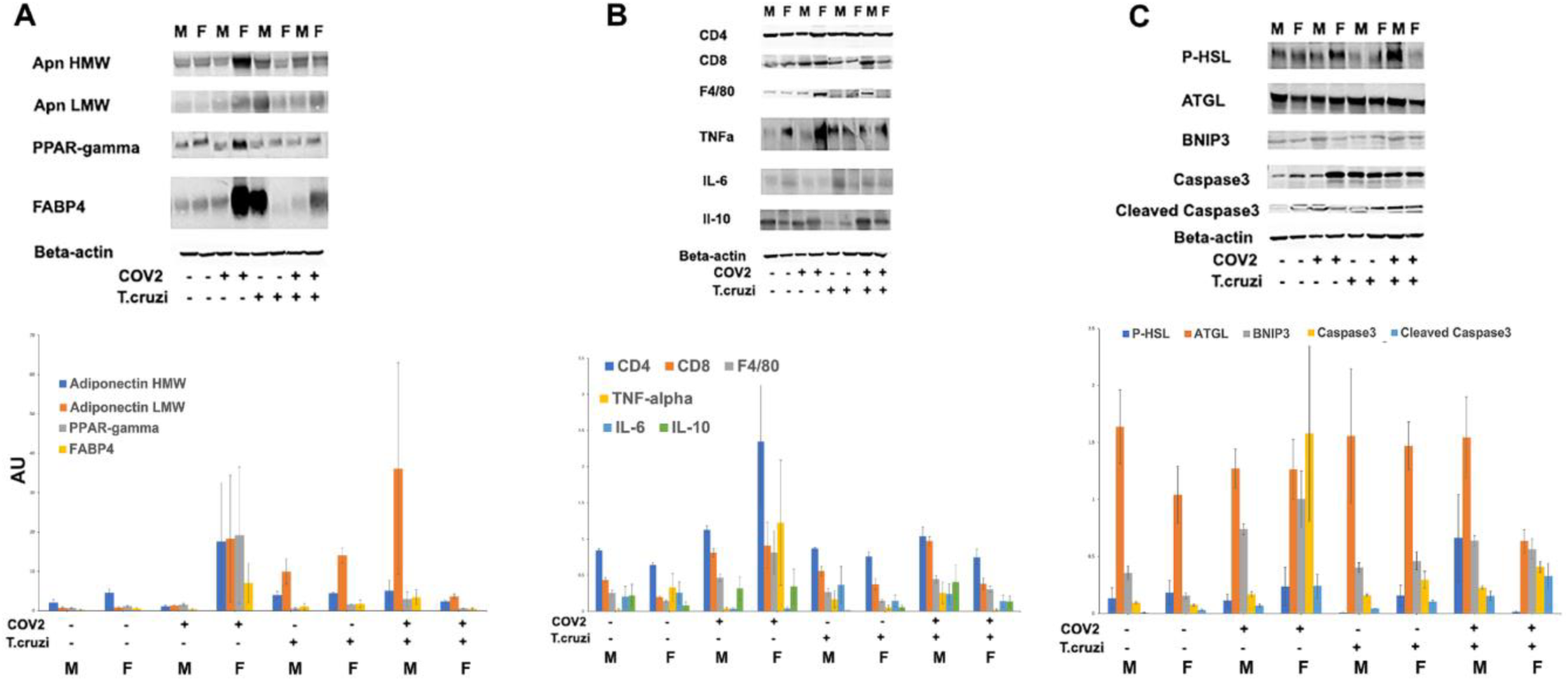
Immunoblot analysis (upper panel) of markers of: A) adipogenesis (HMW ApN, gAd, PPARγ and FABP4); B) Immune cells (CD4, CD8 and F4/80) and inflammatory cytokines (TNF, IFNg); and C) Lipolysis (p-HSL and ATGL), necrosis (BNIP3) and apoptosis (caspase 3 and cleaved caspase3) in the adipose tissue of control and infected (CoV2, *T. cruzi* and coinfected) male (M) and female (F) mice. Bar graphs (lower panels) of the levels of each protein marker normalized to beta-actin for A, B, and C respectively (x axis-arbitrary units). The error bars represent the standard error of the mean. The comparative-fold change in the expression levels of the above protein markers is presented in Table 1B.

**Table 1B.**
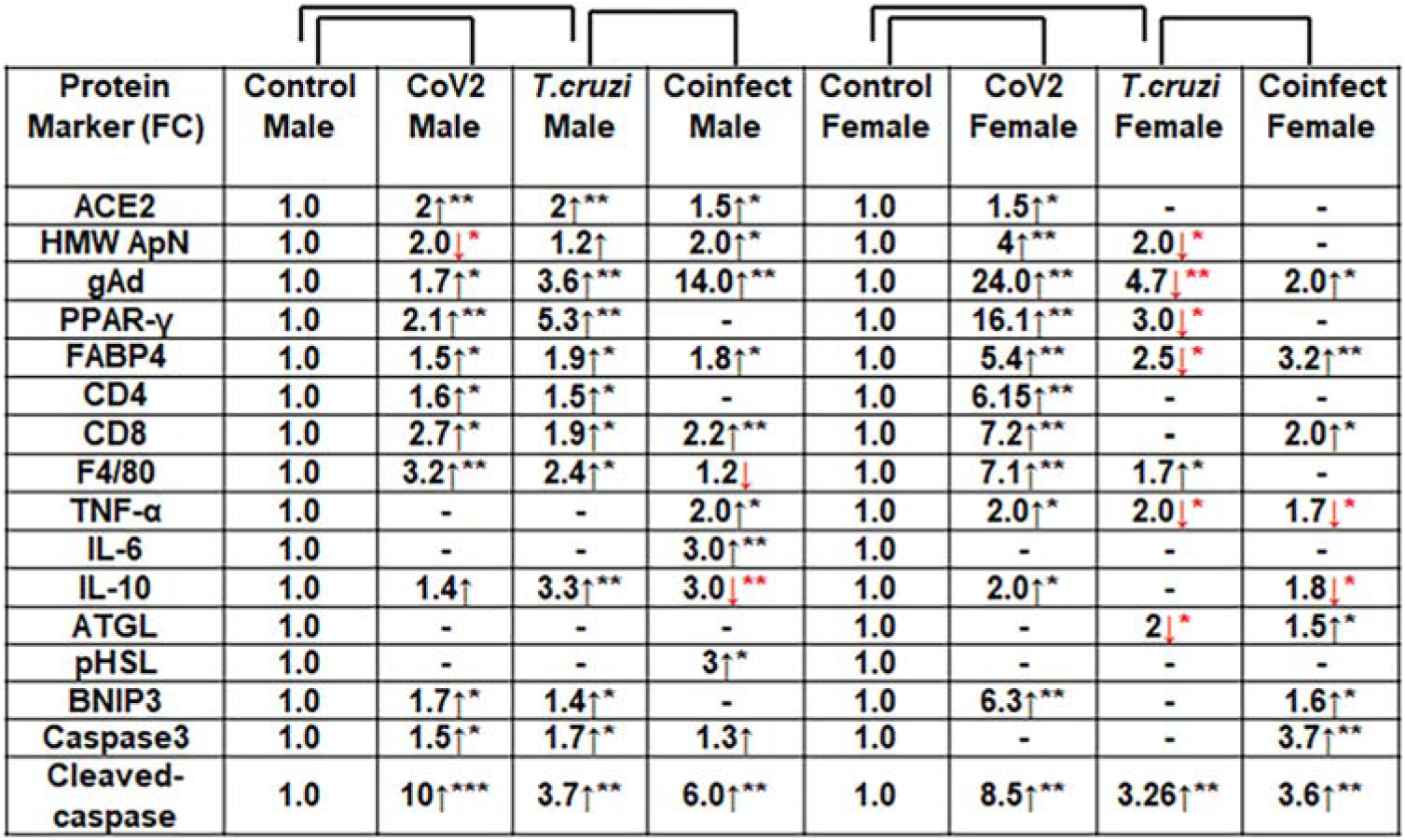
The-fold change of the protein markers (adipogenic, immune and metabolic signaling) levels compared to their sex matched control mice are presented in Table 1B analyzed in adipose tissue (WAT). The-fold changes were analyzed by comparing the protein’s normalized level (GDI or β-actin) in infected groups (CoV2/*T. cruzi*) to that in uninfected (control) mice, for males and females separately. For the coinfected mice, since the baseline is *T. cruzi* infection, the-fold change was calculated for coinfected mice relative to *T. cruzi* infected mice (for males and females separately). The increase and decrease in the comparative-fold change are presented by an upward or downward arrow, respectively (* p ≤ 0.05, ** p ≤ 0.01 and *** p ≤ 0.001 represents the significance). N=3-4/group.

We also analyzed the levels of other adipogenic factors, such as PPARγ and FABP4, in WAT of CoV2 and coinfected mice (Fig. 4A). Similar to the changes in adiponectin levels, the levels of PPARγ and FABP4 significantly increased (16.1-fold and 5.4-fold, respectively) in female COV2 infected mice compared to female control mice and the fold increases were significantly greater in female COV2 infected mice compared to male CoV2 infected mice. *T. cruzi* infection increased the levels of only FABP4 in both male and female mice (1.8- and 3.2-fold, respectively) compared to the respective sex matched control mice (Fig. 4A). Interestingly, CoV2 infection *T. cruzi* infected mice significantly increased the levels of both PPARγ and FABP4 in males (5.3-fold and 1.9-fold, respectively), but significantly decreased both of them in females (3-fold and 2.5-fold, respectively) compared to sex matched *T. cruzi* infected mice (Table 1B). These results demonstrated that adipogenic signaling is increased in female CoV2 infected mice and male coinfected mice compared to their respective sex matched infection controls.

### CoV2 infection and coinfection differently alter immune signaling in WAT in male and female mice

Immunoblot analysis in WAT lysates demonstrated significant differences in the protein levels of WAT immune cells (CD4, CD8, and macrophage marker F4/80) and inflammatory markers (TNFα, IL-6 and IL-10) between the sexes and infections (Fig. 4B, Table 1B). Uninfected female mice showed significantly lower levels of resident CD4 and CD8 cells (1.3-fold and 3.6-fold, respectively) and increased F4/80 levels (1.5-fold) compared to uninfected male mice (Fig. 4B). CoV2 infection increased the infiltration of CD4, CD8 and F4/80 in WAT in both males (1.6-, 2.7- and 3.2-fold, respectively) and females (6.15-, 7.2- and 7.11-fold, respectively) compared to their respective sex matched uninfected mice. However, the levels of immune cells (CD4, CD8 and macrophages) were significantly increased in WAT in female CoV2 mice compared to male CoV2 mice. *T. cruzi* infection increased only CD8 levels in WAT in both male and female mice (2.2-fold and 2-fold, respectively) compared to sex matched control mice. The coinfection significantly increased CD4, CD8 and F4/80 levels (1.5, 1.9 and 2.4-fold, respectively) in WAT in male coinfected mice, whereas only the levels of F4/80 significantly increased (1.7-fold) in female coinfected mice compared to sex matched *T. cruzi* infected mice (Fig. 4B).

The levels of proinflammatory TNFα were significantly higher (2-fold) in WAT in female uninfected mice compared to male uninfected mice (Fig. 4B). CoV2 infection further increased (2-fold) the levels of TNFα in female mice. No significant change in the levels of IL-6 was observed in either male or female CoV2 infected mice compared to the sex matched control groups. *T. cruzi* infection increased the levels of TNFα and IL-6 in WAT in male mice (2- and 3-fold, respectively) and TNFα decreased significantly in female (1.7-fold) mice compared to sex matched control mice (Fig. 4B, Table 1B). CoV2 infection further decreased (2-fold) the levels of TNFα in *T. cruzi* infected female mice (coinfected state) compared to *T. cruzi* infected female mice. The levels of anti-inflammatory IL-10 significantly increased in WAT of males and females (1.4- and 2-fold, respectively) in CoV2 infected mice compared to sex matched control mice. The levels of IL-10 decreased in WAT of male and female (3- and 1.8-fold, respectively) in *T. cruzi* infected mice compared to sex matched control mice. However, the levels of IL-10 significantly increased (3.3-fold) only in WAT of male coinfected mice compared to male *T. cruzi* infected mice, whereas no change was observed in female coinfected mice (Fig. 4B). These data demonstrated that CoV2 infection induces stronger WAT proinflammatory signaling in females compared to males, but that *T. cruzi* coinfection provokes stronger WAT proinflammatory signaling in males compared to females (Table 1B).

### CoV2 infection and coinfection cause different types of cell death (apoptosis vs necrosis) in WAT of male and female mice

Histological analysis demonstrated significant loss of adipocytes in CoV2, *T. cruzi* and coinfected mice compared to their respective control groups (Supplemental Fig. 2). We analyzed whether the cause for the loss of adipocytes was due to apoptosis or necrosis by quantitating the protein levels of cleaved caspase 3 and BNIP3, respectively, in WAT (Fig. 4C). In WAT of uninfected female mice, the levels of cleaved caspase 3 were significantly higher (3.7-fold) and the levels of BNIP3 significantly lower (1.8-fold) compared to uninfected male mice, which suggest that in WAT of female mice the process of cell death is predominantly due to apoptosis. CoV2 infection increased the levels of cleaved caspase 3 and BNIP3 in both males (10- and 1.7-fold, respectively) and females (8.5- and 6.3-fold, respectively) compared to sex matched control mice (Fig. 4C, Table 1B). However, the levels of necrotic cell death were greater in female WAT compared to male WAT in CoV2 infected mice (Table 1B). *T. cruzi* infection increased the levels of cleaved caspase 3 in WAT of both males and females (6- and 3.6-fold, respectively) and increased BNIP3 only in females (1.6-fold) compared to sex matched control mice. Coinfection further increased the levels of cleaved caspase 3 in WAT of both males and females (3.7- and 3.3-fold, respectively) and increased BNIP3 only in males (1.4-fold) compared to sex matched *T. cruzi* infected mice. These data indicated that during CoV2 infection adipose tissue is predominantly lost via apoptotic cell death and that necrotic WAT cell death is greater in female mice compared to male mice. In contrast, in coinfected mice, although apoptotic cell signaling also predominates in both male and female mice, necrotic signaling is higher in male mice compared to female mice.

### Sex dependent morphological changes in the hearts of mice infected with CoV2, *T. cruzi* and coinfection

We have shown that CoV2 infects and persists in the hearts of intra-nasally infected mice (Fig. 1B). Histological analysis of the hearts was performed using H&E and Masson-trichrome stained sections as described in Materials and Methods. Microscopic analysis of the heart sections of CoV2 infected mice demonstrated the presence of infiltrated immune cells, increased accumulation of lipid droplets in the capillaries, enlarged cardiomyocyte nucleus, and fibrosis compared to control mice (Fig. 5A and 5B). The H&E sections showed significantly reduced cytoplasmic coloration in LV in female *T. cruzi* infected and coinfected mice compared to their sex matched counterparts (Fig. 5A). RV in *T. cruzi* infected mice showed increased fibrosis compared to control mice. Coinfected mice showed significantly elevated fibrosis compared to *T. cruzi* infected mice (Fig. 5B). The levels of accumulated lipid droplets in the capillaries, infiltrated immune cells and fibrosis in RV in male coinfected mice were significantly greater compared to female coinfected mice (Fig. 5B) (Supplemental Fig. 3).

**Figure 5.**
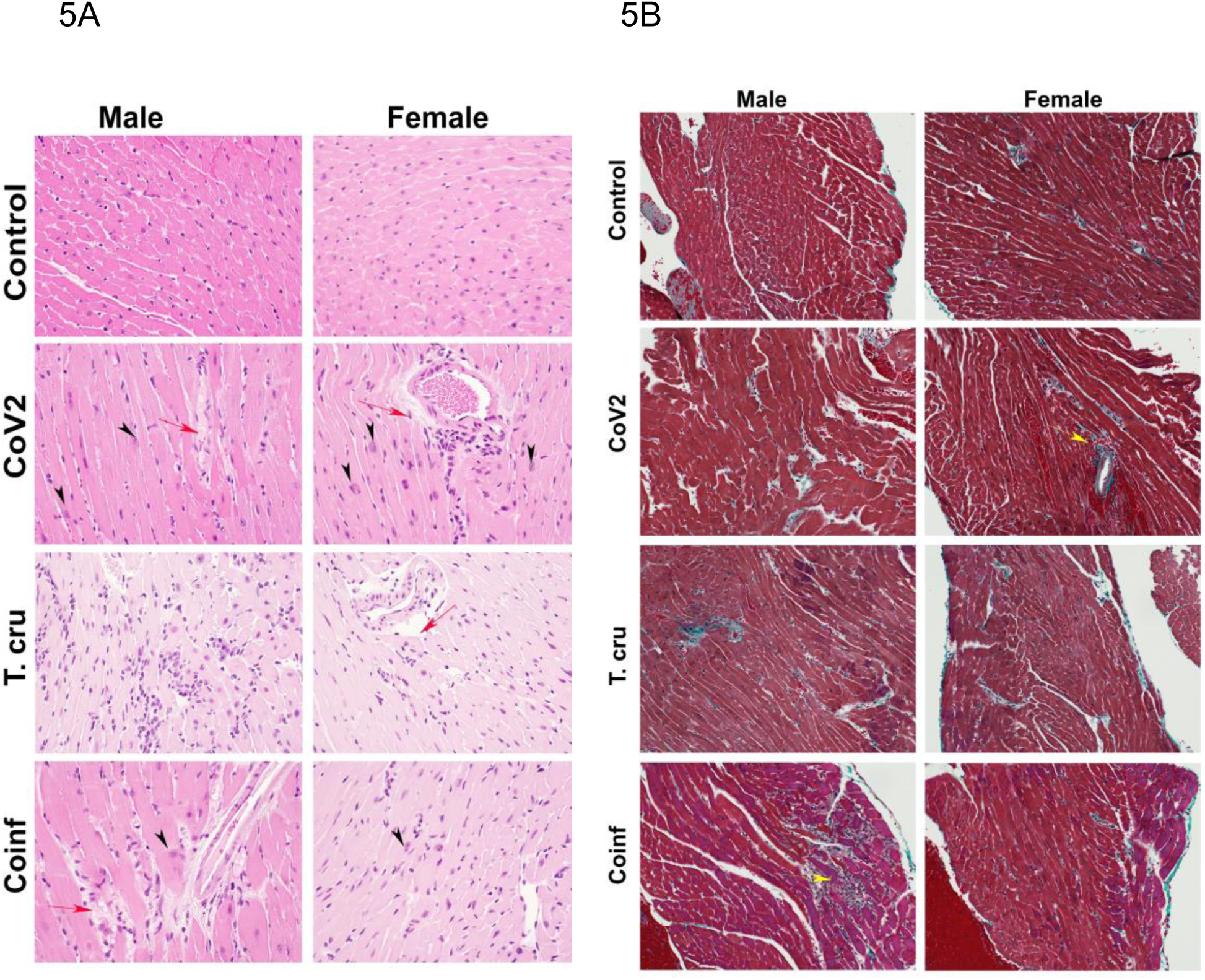
Histology of the myocardium of hACE2 mice infected with CoV2/T. cruzi and coinfected mice (n = 3-4, minimum five images/section were analyzed). (A) H&E staining showed significantly increased lipid droplets in the capillaries (red arrow) and enlarged cardiomyocyte nucleus (black arrowhead) in the left ventricles in CoV2 infected mice relative to uninfected mice and in coinfected mice hearts compared to the hearts of *T. cruzi* infected mice (20x magnification). (B) Masson-trichrome staining showed significantly more fibrosis and damage (immune cells – yellow arrows) in the right ventricles (RV) of infected/coinfected mice compared to uninfected hACE2 mice (20x magnification). (Additional images are presented in supplemental Fig. 3).

We performed the morphometric analysis of the hearts as described in Materials and Methods section. The thickness of the left ventricular wall (LVW), right ventricular wall (RVW) and septal wall (SW) differed between males and females and infected and coinfected mice compared to sex matched control mice (Supplemental Table 1). LVW thickness significantly decreased in female CoV2 and *T. cruzi* infected mice compared to female control mice; however, no significant difference was observed in female coinfected mice. LVW thickness in male CoV2/*T. cruzi* infected and coinfected mice showed no significant differences compared to male control mice. RVW thickness significantly decreased in female control mice compared to male control mice, and was further decreased in female coinfected mice. Interestingly, the thickness of RVW was significantly reduced in male coinfected mice compared to male *T. cruzi* infected mice, which was not observed for female coinfected and *T. cruzi* infected mice. SW thickness increased in female CoV2 mice and was inversely proportional to the decreased LVW thickness compared to female control mice.

### CoV2 infection alters cardiac adiponectin (C-ApN) levels and adiponectin (ApN) signaling in the hearts in coinfected mice

We detected no change in parasite load in the hearts between *T. cruzi* and coinfected mice (data not shown); however, we observed significant heart morphological changes, including accumulation of lipid droplets (Fig. 5). Previously we showed a strong correlation between C-ApN levels and progression of cardiomyopathy during CD, wherein elevated levels of C-ApN were associated with mortality due to cardiac dilation [20]. Here, we used native gel electrophoresis to quantitate HMW-ApN and SDS-PAGE to quantitate the levels of globular ApN (gAd) to determine whether the distribution pattern of ApN in the hearts differed between the sexes and infections (Fig. 6A). The levels of C-HMW ApN did not change, but gAd significantly decreased in uninfected female mice compared to uninfected male mice (Fig. 6A). The-fold changes in the levels of C-HMW Apn and C-gAd in the mice (males and females) during CoV2, *T. cruzi* and coinfection compared to their respective controls are presented in Table 1C. CoV2 and *T. cruzi* infections significantly increased C-HMW ApN levels (1.6- and 4.3-fold, respectively) and significantly decreased gAd (2.5- and 5.0-fold, respectively) in infected male mice compared to uninfected male mice. In females, *T. cruzi* infection significantly increased C-HMW ApN (2.9-fold), whereas both CoV2 and *T. cruzi* infections significantly increased gAd (5.6- and 1.8-fold, respectively) levels compared to uninfected female mice. Although the levels of HMW and gAd differed between the males and females during CoV2 and *T. cruzi* infections, the levels of HMW ApN and gAd showed a similar trend between males and females during coinfection. The levels of HMW decreased in the hearts of coinfected mice, whereas the levels of gAd significantly increased in the hearts of both males (30-fold) and females (23.8-fold) in coinfected mice compared to the sex matched *T. cruzi* infected mice. The data indicate that in CoV2/*T*.*cruzi*/coinfection group males had higher levels of C-HMW ApN, whereas infected females had higher levels of C-gAd compared to their counterparts of the opposite sex (Fig. 6A).

**Figure 6.**
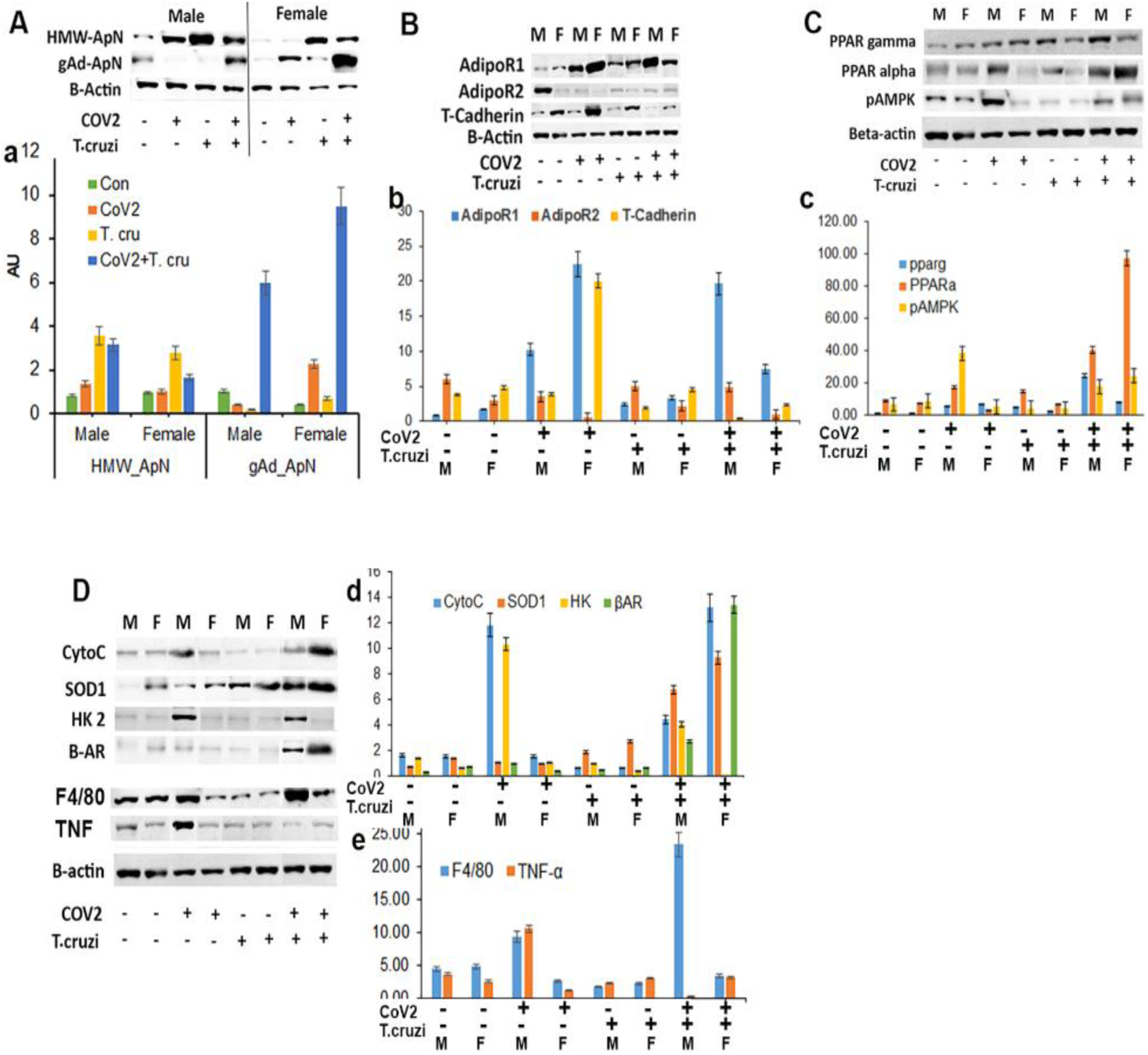
Immunoblot analysis (upper panel) of markers of :(A) adiponectin (HMW ApN and gAd); (B) ApN receptors (Adipo R1, R2 and T-cadherin; (C) lipid metabolism (PPARα and PPARγ) and energy sensor (pAMPK) and (D) mitochondrial markers (Cytochrome C and superoxide dismutase (SOD)), metabolism (hexokinase 2 and adrenergic receptors), and inflammatory markers (F4/80 and TNFα) in the hearts of control and infected (CoV2, *T. cruzi* and coinfected) male (M) and female (F) mice. Bar graphs (lower panels) of the levels of each protein marker normalized to either GDI or beta-actin for A, B, C, and D, respectively (x axis-arbitrary units). The error bars represent the standard error of the mean. The comparative-fold change in the expression levels of the above protein markers is presented in Table 1C.

**Table 1C.**
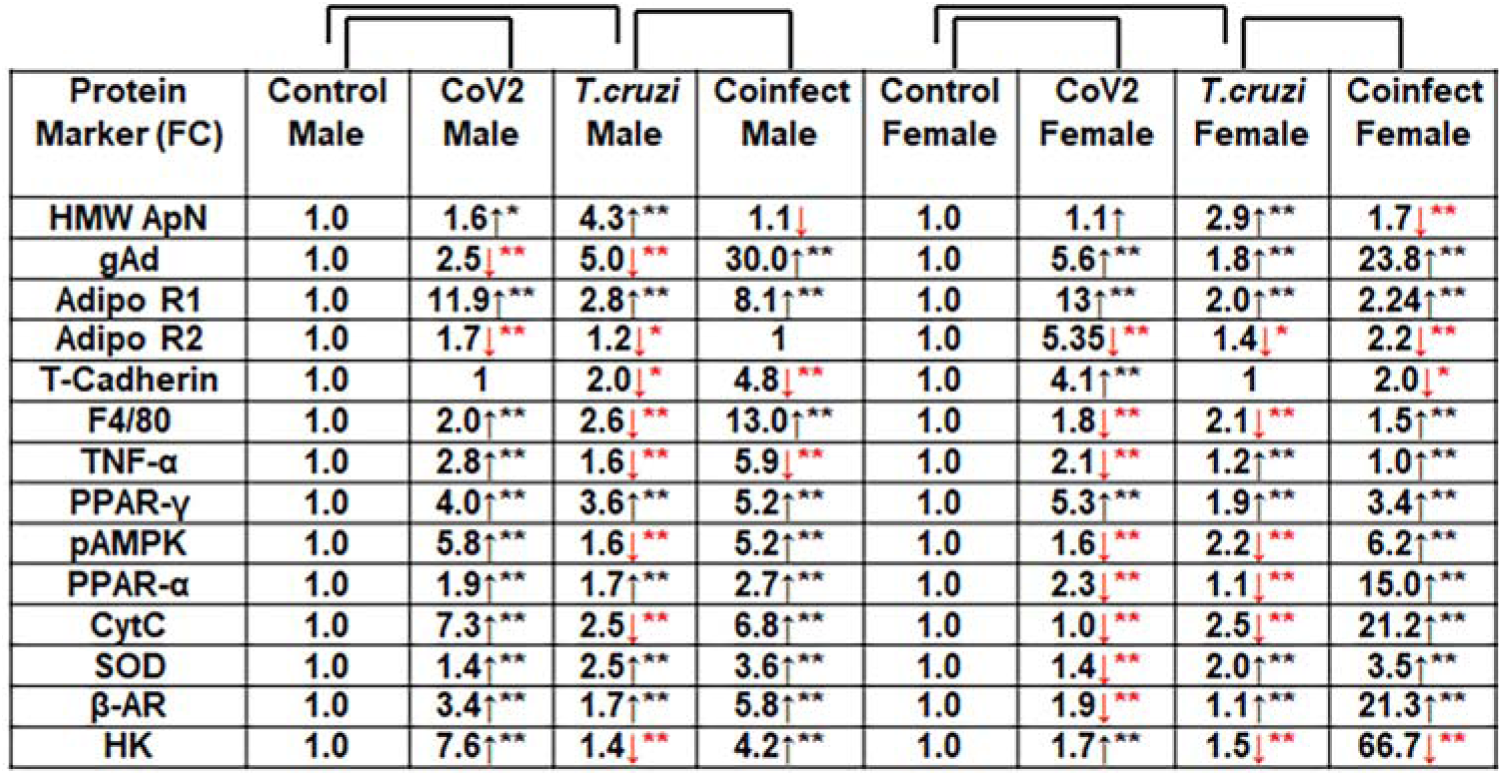
The-fold change of the protein markers (adipogenic, immune and metabolic signaling) levels compared to their sex matched control mice are presented in Table 1B analyzed in the heart. The-fold changes were analyzed by comparing the protein’s normalized level (GDI or β-actin) in infected groups (CoV2/*T. cruzi*) to that in uninfected (control) mice, for males and females separately. For the coinfected mice, since the baseline is *T. cruzi* infection, the-fold change was calculated for coinfected mice relative to *T. cruzi* infected mice (for males and females separately). The increase and decrease in the comparative-fold change are presented by an upward or downward arrow, respectively (* p ≤ 0.05, ** p ≤ 0.01 and *** p ≤ 0.001 represents the significance). N=4/group.

The levels of AdipoR1, AdipoR2 and T-cadherin were significantly altered in the hearts between the sexes and infections (Fig. 6B, Table 1C). In particular, the levels of R1 significantly increased in CoV2 and *T. cruzi* infected mice compared to their sex matched control mice (Fig. 6B). AdipoR1 significantly increased (2-fold) in the hearts of coinfected male mice compared to coinfected female mice (Fig. 6B). The levels of AdipoR2 significantly decreased in infected and coinfected mice compared to their respective sex matched control mice. The levels of AdipoR2 in the hearts in female mice were significantly decreased compared to their male counterparts (Fig. 6B). The levels of T-cadherin significantly increased (4-fold) in female CoV2 mice compared to male coV2 mice (Fig. 6B); however, coinfected mice showed significantly decreased levels of T-cadherin in both male and female mice compared to sex matched *T. cruzi* infected mice (Table 1C).

### CoV2 infection differently alters cardiac lipid and glucose metabolism in the hearts in coinfected male and female mice

ApN regulates lipid (via PPARs) and glucose (AMPK/glycolysis) metabolism [30-33]. We analyzed the levels of PPARs in the hearts as markers of lipid metabolism (lipid oxidation (PPARα), lipogenesis (PPARγ)) and AMPK/pAMPK and hexokinase II (HK) as markers of glucose metabolism (Fig. 6C, Table 1C). Western blotting analysis of PPARs demonstrated significantly increased PPARγ in the hearts of CoV2 and *T. cruzi* infected mice compared to control mice both in males and females. CoV2 infection further significantly elevated the levels of PPARγ in coinfected males (5.2-fold) and coinfected females (3.4-fold) compared to *T. cruzi* infected mice. *T. cruzi* infected and coinfected male mice displayed greater levels of PPARγ compared to their respective female counterparts, suggesting increased adipogenic/lipogenic signaling in males compared to females. The levels of PPARα significantly increased in male CoV2 and *T. cruzi* infected mice (1.9- and 1.7-fold, respectively) and significantly decreased in female CoV2 and *T. cruzi* infected mice (2.3- and 1.1-fold, respectively) compared to sex matched control mice (Fig. 6C, Table 1C). Although coinfection significantly increased PPARα in the hearts of both male (2.7-fold) and female (15-fold) mice compared to sex matched *T. cruzi* infected mice, the levels of PPARα in the hearts of coinfected female mice were significantly greater compared to coinfected male mice (Table 1C). The PPARs regulate cardiac beta 1 adrenergic receptor (β1AR), which regulates lipolysis and contractility of ventricular cardiac muscle [34]. CoV2 infection significantly increased β1AR levels in males (3.4-fold) and significantly decreased β1AR in females (1.9-fold) compared to sex matched control mice (Fig. 6D). However, coinfection significantly increased β1AR levels in the hearts in both male (5.8-fold) and female (21.3-fold) mice compared to sex matched *T. cruzi* infected mice.

The levels of pAMPK and HK significantly increased (5.8- and 7.6-fold, respectively) in CoV2 infected male mice and significantly decreased (1.6- and 1.4-fold, respectively) in *T. cruzi* infected male mice compared to control male mice (Fig. 6C and 6D, Table 1C). However, coinfection significantly increased the levels of pAMPK and HK (5.2- and 4.2-fold, respectively) in male mice compared to *T. cruzi* infected male mice, suggesting that CoV2 infection increases AMPK signaling and glycolysis in the hearts of male mice. In female mice, CoV2 and *T. cruzi* infections significantly decreased the levels of pAMPK (1.6- and 2.2-fold, respectively) compared to uninfected control female mice. The levels of HK significantly increased in female CoV2 mice (1.7-fold) and significantly decreased in female *T. cruzi* infected mice (1.5-fold) compared to control female mice. However, although the levels of pAMPK significantly increased (6.2-fold), the levels of HK significantly decreased (66.7-fold) in the hearts of coinfected female mice compared to *T. cruzi* infected female mice, suggesting that the glycolytic pathways may not serve as the main energy resource in the hearts of female coinfected mice (Table 1C).

Alterations in cardiac lipid and glucose metabolism may modify the mitochondrial energy pathway and production of reactive oxygen species [35, 36]. Increased PPARα elevates mitochondrial β oxidation and release of reactive oxygen species (ROS) in the hearts during infection, which can impact the progression of CCM [36, 37]. Therefore, we analyzed the levels of Cytochrome C (CytoC) and superoxide dismutase (SOD) in the hearts by Western blotting (Fig. 6D, Table 1C). CoV2 infection significantly increased CytoC and SOD levels in males (7.3- and 1.4-fold, respectively) and significantly decreased SOD in females (1.4-fold) compared to sex matched control mice. *T. cruzi* infection significantly decreased CytoC and significantly increased SOD in the hearts in both male and female mice compared to sex matched control mice. Coinfection significantly increased CytoC and SOD levels in males (6.8- and 3.6-fold, respectively) and females (21.2- and 3.5-fold, respectively) compared to sex matched *T. cruzi* infected mice. Although the levels of SOD were similar in the hearts of male and female coinfected mice, the levels of CytoC were significantly greater in the hearts of female mice compared to male mice, showing a positive correlation with their respective PPARα levels. Together, these data suggested that lipid catabolism and oxidation is greater in female hearts compared to male hearts in coinfected mice which may prevent the progression of cardiac dilation due to intracellular lipotoxicity [38]. Increased lipogenesis due to increased PPARγ in the hearts of male coinfected mice may likely induce early dilated cardiomyopathy in post-COVID mice.

### Cardiac immune signaling differs between male and female mice during CoV2 and *T. cruzi* infections and coinfection

Because alteration in cardiac metabolism may affect immune signaling, we analyzed the levels of infiltrated macrophages and the levels of proinflammatory TNFα in the hearts by immunoblot analysis (Fig. 6D). The levels of F4/80 and TNFα significantly increased (2.0- and 2.8-fold, respectively) in the hearts of CoV2 infected male mice and significantly decreased (1.8- and 2.1-fold, respectively) in the hearts of CoV2 infected female mice compared to respective sex matched control mice. We also observed significantly reduced levels of macrophages in *T. cruzi* infected mice. Interestingly, the levels of macrophage marker F4/80 significantly increased in male coinfected mice compared to female coinfected mice, but the levels of TNFα significantly decreased in male coinfected mice compared to female coinfected mice, revealing differences in inflammatory signaling between male and female coinfected mice.

## DISCUSSION

Many clinical and in *vivo* studies have examined the effect of comorbidities such as diabetes, asthma, hypertension and cardiac diseases on the pulmonary pathogenesis and susceptibility to CoV2 infection. However, the effects of metabolic and immunologic changes associated with chronic infectious disease on the risk of developing severe COVID have not been extensively investigated and neither have been the post-COVID effects on the manifestation/activation of other infectious diseases. This study examines (i) the effect of changes in the immune and metabolic status due to *T. cruzi* infection during an indeterminate stage on susceptibility to pulmonary CoV2 infection and (ii) the effect of CoV2 infection on the pathogenesis and risk of developing cardiomyopathy in *T. cruzi* infected and uninfected mice. Moreover, this study assesses whether the relationship during *T. cruzi* and CoV2 infections differs between male and female sexes. Specifically, to understand the interplay between *T. cruzi* and CoV2 infections we used transgenic hACE2 mice (males and females) nasally infected with SARS-CoV2 mice pre-infected with *T. cruzi*. Our study revealed that: (a) *T. cruzi* infection alters immune and metabolic status in the lungs but reduces the pulmonary SARS-CoV2 load in coinfected mice compared to CoV2 alone infected mice, (b) CoV2 infection alters immune and metabolic status in the hearts and may increase the risk of developing cardiomyopathy in *T. cruzi* infected mice, and (c) CoV2 persists in adipose tissue, altering adipose tissue physiology, which may regulate pulmonary pathology during CoV2 infection and coinfection. More importantly, our study showed that the impact of CoV2 and *T. cruzi* infections and coinfection is sex dependent: male CoV2 and *T. cruzi* singly infected mice were more susceptible to developing pulmonary disease and cardiac disease, respectively, female coinfected mice were susceptible to developing pulmonary disease, and male coinfected mice were susceptible to developing post-COVID cardiomyopathy.

The histopathology of the lungs in CoV2 infected and coinfected mice correlated with the viral loads in both male and female mice. Many clinical studies indicate that males are more susceptible to CoV2 and *T. cruzi* infections compared to females [18, 19, 39-42]. Our animal data supported these clinical data and showed increased viral loads and pulmonary pathology in male CoV2 mice compared to female CoV2 mice. Out of the three organs we analyzed for viral load, the lungs showed the greatest levels of the virus, followed by WAT and the hearts. Interestingly, even though the viral load in the lungs in CoV2 infected female mice was lower compared to male mice, the viral load in WAT in female mice was significantly higher compared to male mice. Thus, we observed an inverse correlation between the viral loads in the lungs and in the WAT in CoV2 infected male and female mice. These data suggest that WAT may play a major role in regulating pulmonary viral load and pathology during COVID. Indeed, previously we demonstrated that pathogens like *T. cruzi* and *Mycobacterium tuberculosis* persists in WAT and that loss of fat cells increases the risk of disease manifestation. For example, we showed that loss of body fat correlated to increased cardiomyopathy in chronic Chagas disease murine model and increased pulmonary pathology in aerosol infected TB murine model [20, 22]. Our current study suggests that WAT may serve as a reservoir for CoV2, sparing the lungs from the viral burden and infection severity. It is well documented that females have higher body fat content compared to males and the fat distribution pattern differs between the sexes, which constitute one reason why males are more susceptible to pulmonary CoV2 infection.

Our study shows for the first time that the persistence of CoV2 in WAT alters adipose tissue morphology and adipocyte physiology. We showed a significant decrease in the size of lipid droplets and a loss of lipid droplets in WAT of CoV2 infected mice. Both male and female mice demonstrated increased apoptotic and necrotic cell death during CoV2 infection and *T. cruzi* coinfection. However, female CoV2 mice and male coinfected mice demonstrated increased adipogenic signaling compared to their respective sex groups, which suggests that increased adipogenic signaling might promote adipogenesis and reverse the loss of lipid droplets. Adipogenesis in WAT in female CoV2 and coinfected mice may help the virus to persist in WAT and spare the lungs, as discussed above. In addition, the levels of infiltrated immune cells (CD4, CD8 and macrophages) in the lungs likely prevent high viral load in CoV2 infected female mice (compared to CoV2 infected male mice) by increasing pro-inflammatory TNFα levels. However, the levels of TNFα and IFNγ significantly increased in the lungs in both coinfected males and females compared to sex matched *T. cruzi* infected mice, and these levels were significantly higher in coinfected male mice compared to coinfected female mice (Table 1A), which may explain why the viral load is significantly lower in the lungs of coinfected male mice compared to coinfected female mice. In addition, *T. cruzi* infection-induced increase in infiltrated immune cells likely prevented early replication of virus in the lungs in coinfected mice.

Previously, we showed that loss of adipocytes correlates to pulmonary adipogenesis and ApN levels in *M. tuberculosis* infected mice [22]. This connection between metabolism and immune activation prompted us to analyze the levels of ApN, a metabolic and immune regulator, in the lungs in CoV2 and *T. cruzi* infected and coinfected mice. We measured the levels of lung high-molecular weight ApN (L-HMW ApN), a.k.a. its anti-inflammatory/anti-fibrotic/metabolically active form [26, 27] and lung gAd (L-gAd), a.k.a. its pro-inflammatory form [28]. Our data suggest that female mice respond better to infections like CoV2 and *T. cruzi* than male mice by increasing gAd levels and inducing pro-inflammatory signaling in the lungs. Although the levels of L-HMW ApN significantly increased in *T. cruzi* infected mice, during coinfection their levels significantly decreased, whereas the levels of L-gAd increased. It has been shown that macrophage elastases cleave ApN to generate gAd [43]. Our data suggest that during *T. cruzi* infection the infiltrated macrophages might cleave full length ApN to gAd and create a pro-inflammatory environment at the initial stages of CoV2 infection, thus reducing the viral load in the lungs in coinfected mice compared to CoV2 infected mice.

Similar to the changes in the metabolic and immunologic conditions in the lungs during CoV2 and coinfection, we observed altered HMW-ApN and gAd in the hearts, which significantly differed between males and females (Table 1C). These alterations in ApN levels and adipogenic signaling may regulate energy metabolism differently in the hearts of male and female coinfected mice. Our data showed that higher C-gAd levels correlated with increased PPARα, pAMPK, CytoC, and β1AR and decreased HK in female coinfected hearts compared to male coinfected hearts (Table 1C). Although the levels of SOD increased in female coinfected mice in response to the ROS generated during β-oxidation, these levels of SOD are likely not enough to neutralize all the ROS generated compared to male coinfected mice (compared to CytC levels between the sexes). The significant increase in the levels of β-oxidation may increase the LV contractile power [44]. The levels of β1AR correlated to the levels of PPARα in the hearts, suggesting that PPARα-induced fatty acid oxidation may increase β1AR levels, causing elevated contractility of ventricles. Our data suggest that the significant increase in ApN-PPARα induced mitochondrial β-oxidation of lipids in the hearts may be the cause for the reduced heart size (shrunken heart) in female coinfected mice, a condition similar to that observed in Acetyl-coA carboxylase (ACC2, an enzyme involved in fatty acid biosynthesis) mutant mice [45]. On the other hand, significantly increased HMW ApN levels in the hearts correlated with increased PPARγ and HK in male coinfected mice compared to female coinfected mice (Table 1C), which suggests that the hearts of male coinfected mice may utilize energy derived from glycolysis and store lipids in the form of triglycerides (adipogenesis/lipogenesis).

Increased C-HMW ApN and PPARγ elevates vascular dilation [46, 47]. Earlier we have demonstrated a correlation between increased cardiac ApN, lipid accumulation and cardiomyopathy [37]. We also showed a correlation between altered metabolic status and immune status in the hearts. Although the levels of macrophages significantly increased in male coinfected mice compared to female coinfected mice, TNF levels were significantly decreased in males. These data suggested that increased ApN-PPARγ associated signaling in the hearts of male coinfected mice might have induced an anti-inflammatory response by altering macrophage polarization from the M1 to the M2 form. Thus, in coinfected mice, males and females showed different heart phenotypes, which correlated with increased PPARγ and PPARα levels, respectively. Overall, these data suggest that CoV2 infection and coinfection with *T. cruzi* differently affect cardiac metabolic and immune status in male and female mice via host C-ApN levels and signaling. Thus, the C-ApN-PPARα and ApN-PPARγ signaling axes may play major roles in determining the progression and severity of CCM in the context of COVID.

The present study investigated the immediate effect of CoV2 infection on the pathology of the lungs and hearts in CoV2 and *T. cruzi* (coinfection) infected mice, while any potential long-term effects still remain to be explored. Further studies including a greater number of male and female mice at different time stamps are warranted to evaluate the long-term post-COVID effects on the development and progression of Chagas cardiomyopathy.

## CONCLUSION

Our data demonstrated that SARS-CoV2 infects adipocytes, persists in adipose tissue, and causes a loss of adipocytes and lipid droplets. The loss of fat cells correlates to the pulmonary adipogenic signaling via ApN isomers. We showed that the levels of L-HMW and gAd differ between male and female mice during CoV2 infection and coinfection, which may differently regulate inflammation, viral load, and pathology in the lungs in males and females. These findings may underpin the clinical observations that males are more susceptible to COVID than females and suffer greater pulmonary damage. Our study also suggests that the severity of CoV2 infection may be lower in the lungs of *T. cruzi* pre-infected subjects due to increased proinflammatory status in the lungs. However, the risk of developing dilated cardiomyopathy in *T. cruzi* infected males may be greater than females coinfected with CoV2.

## ACKNOWLEDGMENTS

We thank Erika Shor at the Center for Discovery and Innovation, Hackensack University for a critical reading of the manuscript. We also thank Steven Park at the CDI for the managerial support to executing the BSL3 work. This study was supported by grants from the National Institute of Allergy and Infectious Diseases (National Institutes of Health AI150765-01) to Jyothi Nagajyothi.

None of the authors have a conflict of interest.

## Supplemental Figures

**Supplemental Figure 1**. Flow chart of experimental design.

**Supplemental Figure 2**. Histological images of H&E-stained sections of white adipose tissue (WAT) showing the alteration in the morphology of adipocytes and adipose tissue in mice infected with CoV2, *T. cruzi* and coinfection.

**Supplemental Figure 3**. H&E sections (top panel) and Masson trichrome images (bottom panel) of right ventricles (RV) of coinfected male and female mice showing fibrosis (blue and purple), infiltration of immune cells (yellow arrow), accumulation of lipid droplets (red arrow) and enlarged nucleus (green arrowhead) (40X magnified).

**Supplemental Table 1**. Morphometric analysis of the hearts of CoV2, *T. cruzi* infected and coinfected male and female mice. The thickness of the right ventricle wall (RVW), left ventricle wall (LVW) and intra-septal wall (Septal -W) is measured as mentioned in materials and methods and presented in mm. The significance difference in the wall thickness calculated by t-test comparing to the sex matched uninfected mice denoted by “*” (* p ≤ 0.05, ** p ≤ 0.01 and *** p ≤ 0.001). The significance difference in the wall thickness between coinfected and sex matched *T. cruzi* infected mice denoted by “#” (# p ≤ 0.05). (M-male; F-female; Con-control; CoV-CoV2 infected; T.c-*T. cruzi* infected; Coinf-coinfected)

## Notes

### Competing Interest Statement

The authors have declared no competing interest.

