## supplemental files for "Sex differences in cardio-pulmonary pathology of SARS-CoV2 infected and *Trypanosoma cruzi* co-infected mice"

**Supplemental Fig. 1:**
**Experimental design**

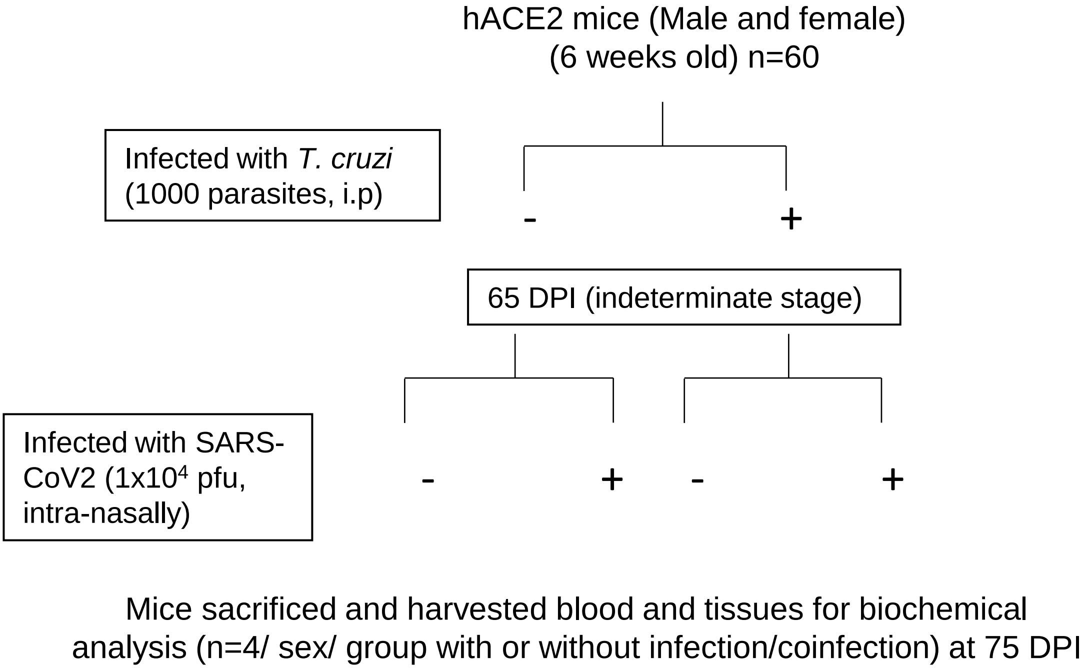

**Supplemental Fig. 2:**

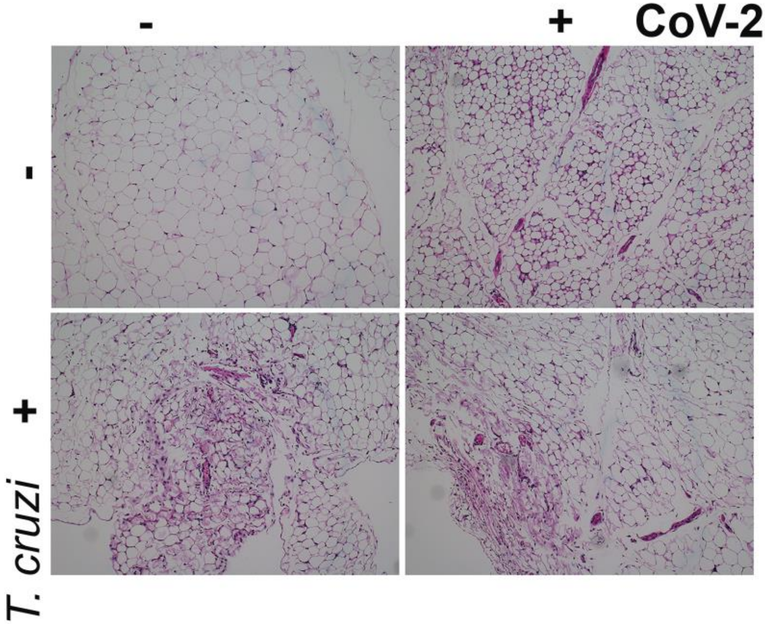

Supplemental Fig. 3

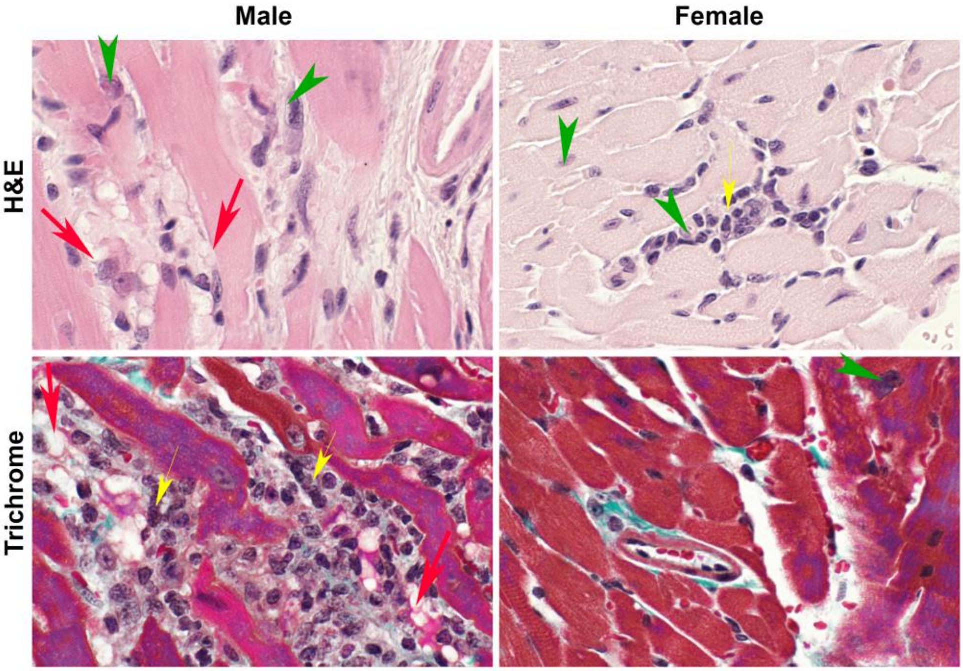

Supplemental Table 1

| | Con_M | Con_F | CoV_M | CoV_F | T.c_M | T.c_F | Coinf_M | Coinf_F |
|---|---|---|---|---|---|---|---|---|
| **RVW** | 1354.5±148 | 1212.4±161 | 1104±228 | 1351±23 | ***<br>2129±159 | 1162±99 | #<br>1285±83 | *<br>927±84 |
| **LVW** | 1694±26 | 1543±46 | 1804±87 | *<br>1383±84 | 1735±19 | *<br>1254±119 | 1850±77 | *<br>1850±31 |
| **Septal W** | 3375±95 | 3909±189 | 3555±81 | *<br>4434±170 | **<br>4001±17.3 | **<br>2828±319 | 3677±165 | *<br>2775±85 |
